# Clustering of loci controlling species differences in male chemical bouquets of sympatric *Heliconius* butterflies

**DOI:** 10.1101/2020.06.03.131557

**Authors:** Kelsey J. R. P. Byers, Kathy Darragh, Sylvia Fernanda Garza, Diana Abondano Almeida, Ian A. Warren, Pasi M. A. Rastas, Richard M. Merrill, Stefan Schulz, W. Owen McMillan, Chris D. Jiggins

## Abstract

The degree to which loci promoting reproductive isolation cluster in the genome – *i.e.* the genetic architecture of reproductive isolation - can influence the tempo and mode of speciation. Tight linkage between these loci can facilitate speciation in the face of gene flow. Pheromones play a role in reproductive isolation in many Lepidoptera species, and the role of endogenously-produced compounds as secondary metabolites decreases the likelihood of pleiotropy associated with many barrier loci. *Heliconius* butterflies use male sex pheromones to both court females (aphrodisiac wing pheromones) and ward off male courtship (male-transferred anti-aphrodisiac genital pheromones), and it is likely that these compounds play a role in reproductive isolation between *Heliconius* species. Using a set of backcross hybrids between *H. melpomene* and *H. cydno*, we investigated the genetic architecture of putative male pheromone compound production. We found a set of 40 significant quantitative trait loci (QTL) representing 33 potential pheromone compounds. QTL clustered significantly on two chromosomes, chromosome 8 for genital compounds and chromosome 20 for wing compounds, and chromosome 20 was enriched for potential pheromone biosynthesis genes. There was minimal overlap between pheromone QTL and known QTL for mate choice and color pattern. Nonetheless, we did detect linkage between a QTL for wing androconial area and *optix*, a color pattern locus known to play a role in reproductive isolation in these species. This tight clustering of putative pheromone loci might contribute to coincident reproductive isolating barriers, facilitating speciation despite ongoing gene flow.

## Introduction

The genetic architecture of population differences can profoundly affect the evolution and maintenance of new species, especially in the face of ongoing gene flow. For example, large effect loci are expected to facilitate speciation (Via, 2012, Merrill et al., 2019) because they are less likely to be lost to drift (Kimura, 1983) and a single large effect locus can lead to substantial reproductive isolation on its own (Bradshaw & Schemske, 2003). Similarly, tight physical linkage of barrier loci, *i.e.* those that contribute to reproductive isolation, will promote speciation by impeding the breakdown of genetic associations between traits that isolate emerging species (Felsenstein, 1981, Smadja & Butlin, 2011). As a result, there is considerable interest in finding the genetic basis of traits that contribute to reproductive isolation and understanding both the effect size and distribution of loci across the genome.

Physical linkage between barrier traits has been detected in a variety of taxa, including pea aphids (Hawthorne & Via, 2001), Laupala crickets (Wiley et al., 2011), *Heliconius* butterflies (Merrill et al., 2019), Ficedula flycatchers (Sæther et al., 2007), stickleback fish (Bay et al., 2017), and *Aquilegia* columbines (Hodges et al., 2002). Barrier traits can include sex pheromones, chemical signals that mediate intraspecific communication important for mating. Due to their critical role in mate attraction, and their ability to convey information about species identity and male quality, pheromones can be important for establishing and maintaining reproductive isolation through relatively simple changes in chemical bouquets (Smadja & Butlin, 2009). As products of secondary metabolism (*i.e.* products not required for survival), alterations in endogenously-produced or -modified pheromones also have the potential to be both relatively simple at the molecular level and, if such changes are relatively late in the pathway, likely avoid major pleiotropic consequences.

Some of the best studied sex pheromones involved in speciation are those of female Lepidoptera, and in some cases the genetic basis of variation in these pheromones is now well known. Both desaturases and fatty acyl-CoA reductases appear to be commonly involved in pheromone biosynthetic variation. Desaturases introduce double or triple bonds between carbon molecules in pheromone components, for example turning alkanes (less common as pheromone components) into alkenes (a common component of lepidopteran pheromones). Fatty acyl-CoA reductases, or FARs, turn fatty acids into fatty alcohols, for example turning octadecanoic acid into octadecanol. In Ctenopseustis, for example, differential regulation of a desaturase drives differences in ratios of sex pheromone components between species (Albre et al., 2012). Desaturases are also important in *Ostrinia* (Roelofs et al., 2002, Sakai et al., 2009, Fujii et al., 2015) and *Helicoverpa* (Li et al., 2015, 2017). Fatty acyl-CoA reductases are responsible for species-specific pheromone alterations in *Ostrinia nubialis* strains (Lassance et al., 2010, 2013). Within single species, desaturases and FARs have received the most attention, being functionally characterized in a variety of Lepidoptera including *Yponomeuta* (Liénard et al 2010), *Agrotis* (Ding & Löfstedt, 2015), *Manduca* (Buček et al., 2015), *Bombyx* (Moto et al., 2004), and *Antheraea* (Wang et al., 2010). In some cases, variation in these female pheromones have also been shown to contribute to reproductive isolation (e.g. Liebherr & Roelofs, 1975, Wu et al., 1999, Emelianov et al., 2001; reviewed in Smadja & Butlin, 2009).

In contrast, male pheromones in Lepidoptera appear more varied, with a diverse array of compound classes (Conner & Iyengar, 2016, Löfstedt et al., 2016). Many male Lepidopteran pheromones resemble plant compounds and are thought to be diet-derived, especially terpenoids and pyrrolizine alkaloids, the latter of which can be derived from either larval or adult feeding (Conner & Iyengar, 2016). In comparison to female pheromones, the genetic basis of variation in male sex pheromones has received less attention. In male *Bicyclus anynana*, a desaturase (*Ban-Δ11*) and two FARs, *Ban-wFAR1 and Ban-wFAR2*) produce pheromone components and precursors (Liénard et al., 2014), and in the moth *Ostrinia nubialis*, males use the same Δ11-desaturase and Δ14-desaturase as females to produce pheromones (Lassance & Löfstedt, 2009). However, the role of these genes in mediating differences between species remains unclear.

Speciation is typically thought to rely on the accumulation of multiple reproductive barriers, involving divergence in many different traits (Butlin & Smadja, 2017). In order to understand the extent of genetic linkage that underlies species differences it will therefore be useful to study taxa in which multiple different traits can be mapped. *Heliconius* butterflies have been well studied in the context of speciation (Bates, 1861, Merrill et al., 2015, Jiggins, 2017), and in particular we know a great deal about the species complex of *Heliconius melpomene* and its sister lineage *Heliconius cydno/timareta*. Genetic loci underlying wing pattern and mate preference have been mapped in this group (Merrill et al., 2010, Jiggins, 2017, Merrill et al., 2019). The additional role of chemical signaling in reproductive isolation has long been suspected (Jiggins, 2008), but only recently studied in any detail. Chemical profiles of the male wing androconia (patches of specialist scales on male wings that release pheromones) and genital regions (thought to act as aphrodisiacs and anti-aphrodisiacs, respectively) differ between species (Estrada et al., 2011, Mann et al., 2017, Darragh et al., 2020), and are important for mate choice, including altering behavior towards con- and heterospecific individuals (Mérot et al., 2015, Darragh et al., 2017, González-Rojas et al., 2020). Genital pheromones play a more complex role in mate choice in *Heliconius*, as they are transferred by males to females during mating and subsequently decrease advances by other males (Gilbert, 1976, Schulz et al., 2007, Estrada, 2009).

One challenge of studying the entire chemical profile is determining which (combination of) compounds are behaviorally active. The chemical bouquets of wings and genitals are complex, often consisting of 30-70 compounds (Mann et al., 2017, Darragh et al., 2017, 2020, Byers et al., 2020, this study), but it remains unknown how many of these are important for signaling. A single wing compound in *H. melpomene*, octadecanal, is known to be biologically active in *H. melpomene* and *H. cydno* (Byers et al., 2020), while the genital compound (*E*)-β-ocimene acts as the main anti-aphrodisiac in *H. melpomene* (Schulz et al., 2007). In addition, one bioactive genital compound in *H. cydno* (hexyl 3-methylbutyrate, Estrada, 2009) is known. It seems likely that additional compounds in the complex bouquet (30-70 compounds) produced by each of these species are biologically active, but this remains to be tested. The simple genetic control of pheromone components in other systems and their role as secondary metabolism products suggests that they can be altered relatively simply without major pleiotropic consequences to essential organismal functions, as has been seen in some moth species (Groot et al., 2019), though pleiotropic effects have been seen in *Drosophila* (Bousquet et al., 2012 and Zelle et al., 2020). In *Drosophila* some loci specifically showed pleiotropy of production and perception of pheromones, thought to be rare in Lepidoptera (Haynes, 2016). In combination with their importance in inter- and intraspecific mate choice, such simple genetic control and relatively lower risk of pleiotropy makes them ripe material for adaptation and speciation. We here conduct quantitative trait locus (QTL) analyses for wing androconial area and wing and genital compounds that differ between *Heliconius melpomene* L. and *H. cydno* Doubleday (Figure 1; see also Byers et al., 2020, Darragh et al., 2020). We then investigated the patterns of QTL distribution across the genome to test for clustering of loci across chromosomes.

**Figure 1:**
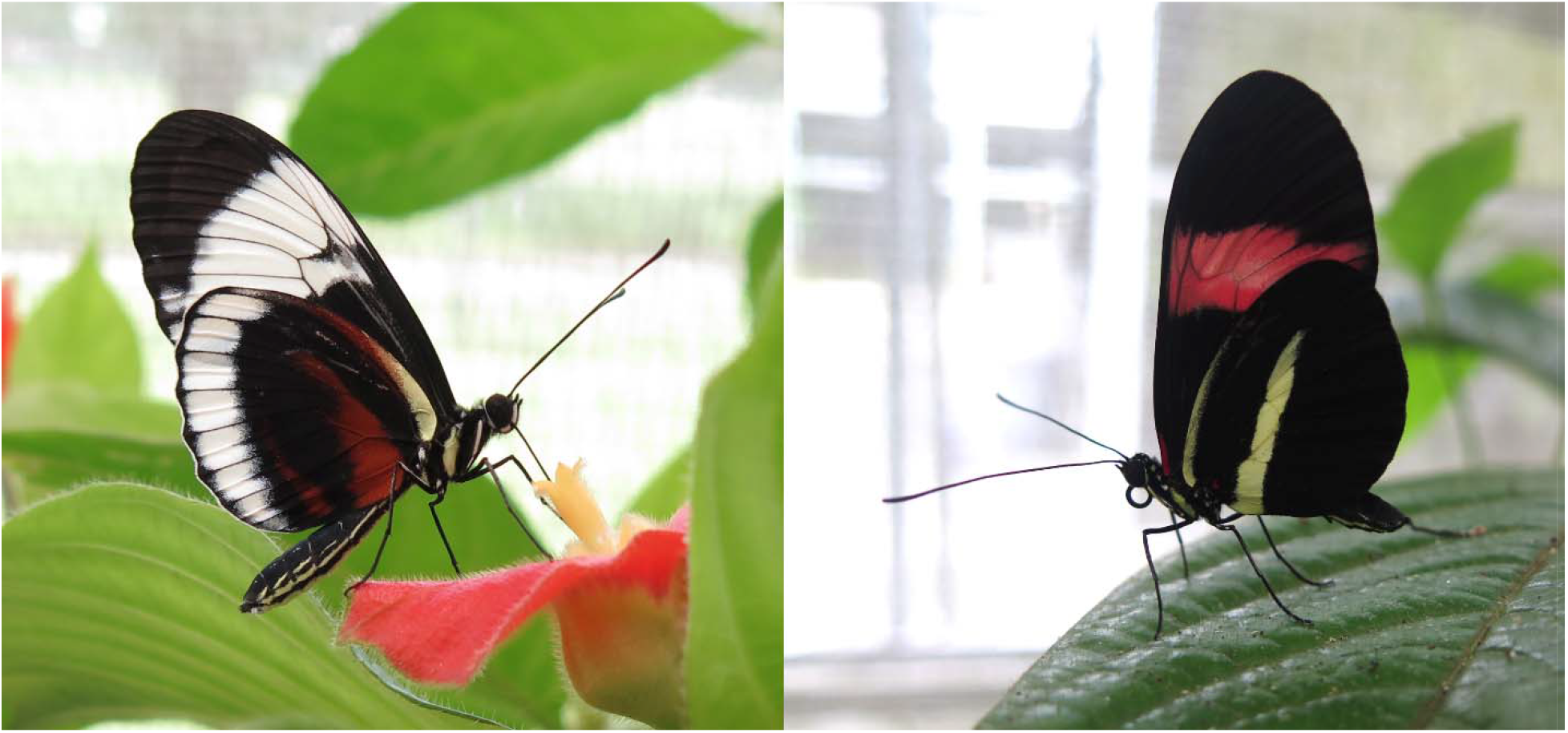
*Heliconius cydno chioneus* (left) and *Heliconius melpomene rosina* (right), the two butterfly species whose putative pheromone components and wing androconial area are mapped in this study.

## Methods

*Heliconius melpomene rosina* and *H. cydno chioneus* (hereafter *H. melpomene* and *H. cydno*) and their interspecific hybrids were reared in outdoor insectaries at the Smithsonian Tropical Research Institute in Gamboa, Panama. Initial stocks were established from wild-caught individuals, and outbred stocks were refreshed with more wild-caught individuals as necessary. Sexes were kept separately and virgin females were kept separate from egg-laying females. To control for host plant effects, all larvae were fed on the same host plant (*Passiflora platyloba* var. *williamsi*), and adults were maintained on a 20% sugar solution and *Gurania eriantha, Psiguria triphylla, Psiguria warscewiczii*, and *Psychotria poeppigiana* as pollen sources. Pheromones were collected from male individuals (both parental species and offspring of mapping crosses) following (Darragh et al. 2017). Wing and genital data from *H. melpomene* and wing data from *H. cydno* have previously been published in (Darragh et al., 2017, Darragh et al., 2019, Byers et al., 2020). Briefly, males of 7-14 days post eclosion had their wings and genitals removed and soaked in dichloromethane plus 1 ng/μL 2-tetradecyl acetate (DCM+IS) for one hour, after which the DCM+IS was removed to a fresh vial and stored at −20°C before gas chromatography-mass spectrometry analysis. Bodies were collected and preserved in 20% dimethyl sulfoxide (DMSO) and 0.25 M EDTA (pH 8.0) and stored at −20 °C for later DNA extraction.

Wing and genital extractions were run on an Agilent 7890B gas chromatograph coupled with an Agilent 5977 mass spectrometer (GC-MS) with electron ionization (Agilent Technologies, California, USA) with an HP-5MS capillary column (30m length, 0.25mm inner diameter) and helium as the carrier gas (1.2 mL/min). Injection used an Agilent ALS 7693 autosampler and was splitless, with an inlet temperature of 250°C. The oven was first held isothermally at 50°C for five minutes, then increased at 5°C/minute to 320°C and again held isothermally for five minutes. Compounds were identified with a custom MS library and quantified using the internal standard area. Only compounds that could be definitively identified were included in the data set. To be included in the mapping experiment, compounds had to be present in at least two thirds of the individuals from one of the parent species, i.e. they had to be “typical” *H. melpomene* or *H. cydno* compounds. A limited set of wing compounds (three methyloctadecanals and two henicosenes from *H. melpomene* and *H. cydno* respectively) were excluded from the wing analysis due to close retention times in the GC-MS data preventing definitive identification of the species phenotype in the mapping individuals. Absolute androconial area and absolute hindwing area were measured in the GNU Image Manipulation Program (GIMP) as pixel counts, then absolute androconial area was divided by hindwing area to produce the percentage of the hindwing taken by the androconial region, the relative androconial area.

Mapping crosses consisted of backcrosses of either an *H. cydno* or *H. melpomene* mother and an F_1_ father, as F_1_ females are normally sterile. Ten families were constructed as backcrosses to *H. melpomene* (representing 89 individuals for the wing phenotype, 81 individuals for the genital phenotype, and 78 individuals for the hindwing relative androconial area phenotype) and fifteen families were constructed as backcrosses to *H. cydno* (127 individuals for the wing, 114 individuals for the genital phenotype, and 124 individuals for the hindwing relative androconial area phenotype) for a total of 216 and 195 hybrid individuals for wing and genital studies respectively. DNA extraction, library preparation, and QTL map construction are detailed in (Byers et al., 2020, Darragh et al., bioRxiv). Briefly, Qiagen DNeasy kits (Qiagen) were used for DNA extraction, and individuals then genotyped either by RAD-seq or low-coverage whole genome sequencing using nextera-based libraries (Picelli et al., 2014, Davey et al., 2017, Merrill et al., 2019). Samples were sequenced by HiSeq 3000 (Illumina) by BGI (China). Linkage mapping was conducted using the standard Lep-MAP3 (LM3) pipeline (Rastas, 2017). The initial linkage groups and marker order were constructed based on the *H. melpomene* genome for 21 chromosomes, as *H. melpomene* and *H. cydno* have highly colinear genomes (Davey et al., 2017). This resulted in a linkage map with 447,818 SNP markers, which was then evenly thinned by a factor of ten to 44,782 markers to ease computation.

To determine which cross direction to use for each compound, we inspected the distribution of the compound phenotype in the parents, F_1_ individuals, and the backcross individuals. Compounds that were present in equal amounts in both parent species (assessed using a Kruskal-Wallis test) were not mapped. Once cross direction was determined, the remaining compounds were log-transformed to approximate normality before being regressed against the linkage map using R/qtl2 (Broman et al., 2018). Due to the family structure present in our crosses, we additionally included a kinship matrix calculated by R/qtl2 using the LOCO (leave one chromosome out) method. Permutation testing was used (with 1000 replicates) to determine QTL threshold. Compounds are likely to be correlated and our data set represents multiple testing of the same mapping crosses, therefore, we used Bonferroni correction separately for each cross direction to establish a second, more conservative, LOD threshold for significance by dividing the threshold by the number of compounds (traits) mapped in the relevant backcross direction. No correction was used for relative androconial area mapping as the trait was mapped alone due to its general lack of correlation with individual volatile compounds. QTL confidence intervals were obtained using bayes_int. As R/qtl2 does not provide a function to calculate the traditional percentage of variance explained by a specific marker, we instead calculated the percentage of the parental difference explained by the genotype at a given marker as a fraction with the difference between the average phenotype of the two genotypes as the numerator and the difference between the average phenotype of the two parental species as the denominator. This can result in values over 100% when more variance is present in the mapping population than the difference between the parent species accounts for.

To identify clustering of QTL, we first calculated the expected number of QTL per chromosome, taking into account the chromosome’s length in cM, then compared that against the observed distribution of QTL per chromosome using a chi-squared test, computing p-values with Monte Carlo permutation with B = 10,000 permutations. Examination of the residuals was used to determine which chromosome(s) displayed clustering, with any residual over 2 considered significant, as in (Erickson et al., 2016). We also repeated these tests taking into account chromosome length in basepairs because the relationship between map length and physical length can differ across chromosomes. The results were unchanged.

As our linkage map is based on whole genome sequencing data, we were able to recover the basepair intervals corresponding to the QTL centimorgan (cM) confidence intervals (defined as the outermost markers at those cM positions) as well as their peaks (defined as the first marker in at that cM position). Lepbase (Challis et al., 2016) was queried to identify genes within the main interval on chromosome 20 for wing fatty acid-derived compound production (identified as the minimum overlapping window of all Bonferroni-significant compound confidence intervals). Genes were searched against the nr (non-redundant) protein database using BLASTp (Altschul et al., 1990) to obtain putative functional annotations, with no specific cutoff used to define a putative annotation; annotations were made when multiple hits with high alignment scores agreed on putative function.

## Results

We first looked at the distribution of compounds in *H. melpomene*, *H. cydno*, and their hybrids. Analysis of the pheromones found a total of 31 compounds in the wings and 68 compounds in the genitals across the two species that were possible to map (i.e. not unknown or overlapping in the gas chromatography-mass spectrometry trace) (Figure 2, Table 1). There was limited overlap in compounds between the wings and genitals, with only the straight-chain alkanes henicosane, docosane, tricosane, pentacosane, and hexacosane (all of which may be part of the normal whole-body cuticular hydrocarbon profile), the unsaturated aldehyde (*Z*)-11-icosenal, and the nitrogenous aromatic benzyl cyanide found in both body regions. A subset of compounds (6 in the wings and 18 in the genitals) were not mapped as they did not differ significantly between the parental species. The rest showed significant differences and thus had adequate variation for QTL mapping to be feasible. Of the wing compounds, 11 showed segregation in backcrosses to *H. cydno* and thus were mapped with those individuals; nine were mapped in backcrosses to *H. melpomene*; and five showed segregation in both backcross directions and thus were mapped separately in backcrosses to each species. For the genital compounds, 19 were mapped in backcrosses to *H. cydno*; 17 in backcrosses to *H. melpomene*; and 14 in both directions.

**Figure 2:**
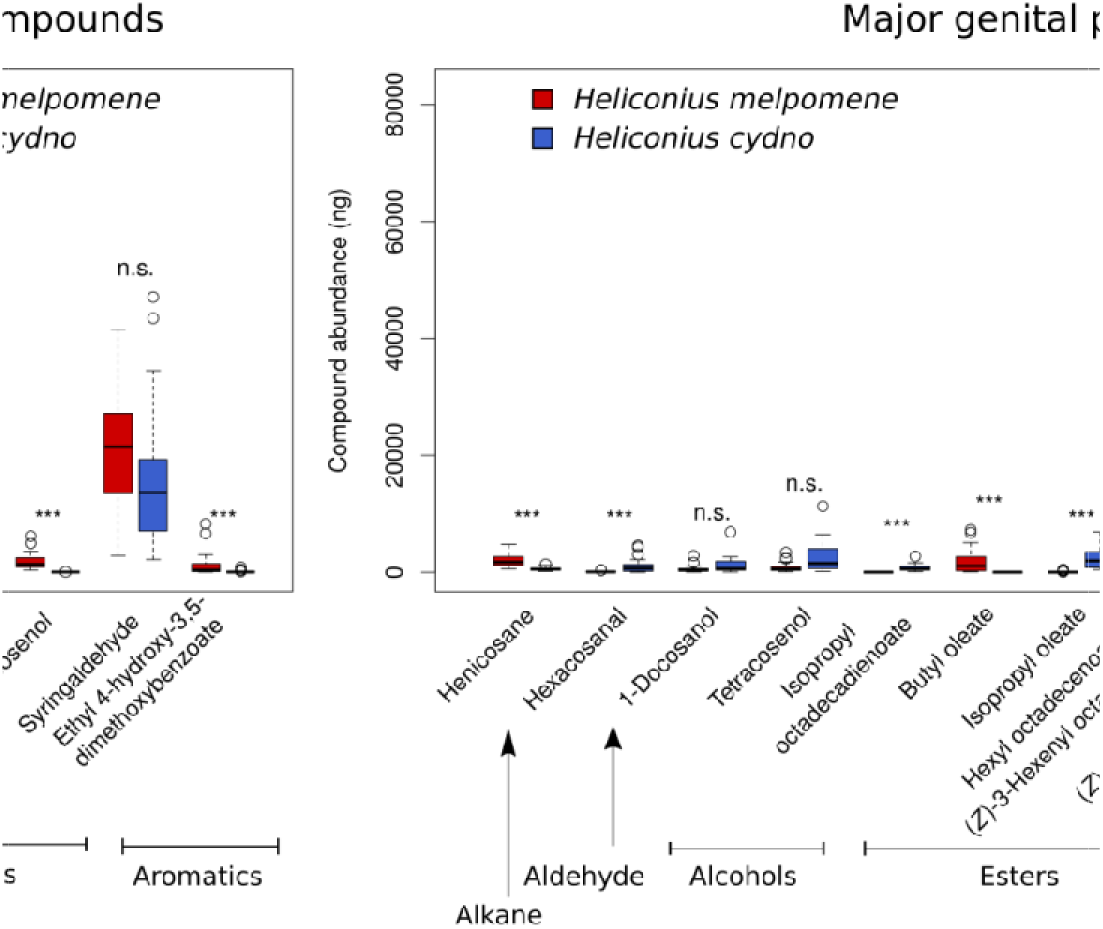
Major compounds found in the wings (left) and genitals (right) of *Heliconius melpomene* and *H. cydno*, arranged by compound type. Only those compounds comprising at least 1% of either species’ pheromone bouquet are included. n.s., not significant; ***: p < 0.005. Boxplots: line is the median, box outline the first and third quartiles, whiskers the most extreme point no more than 1.5 times the interquartile range. [image rotated for reviewer ease of viewing]

**Table 1:**
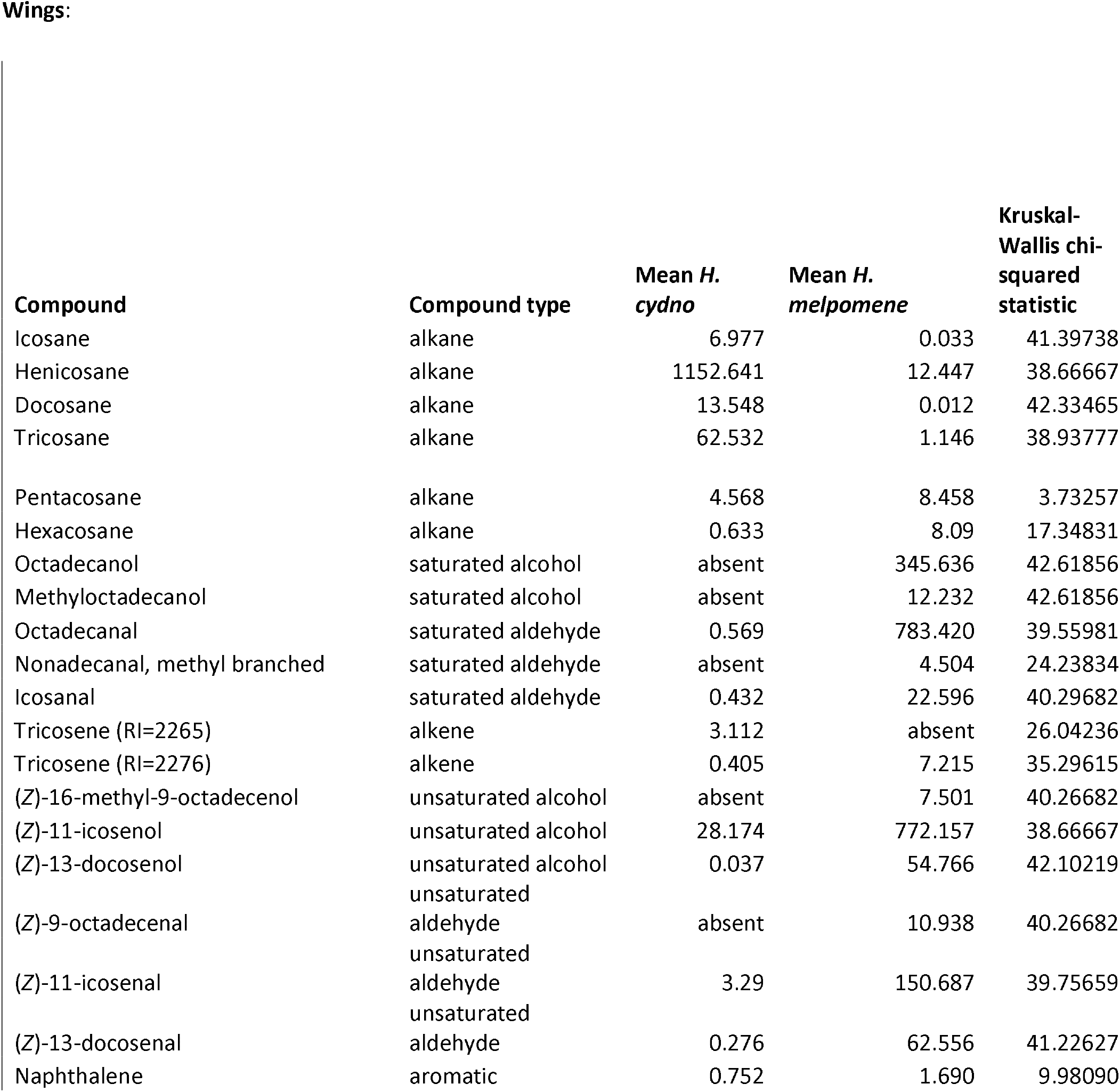
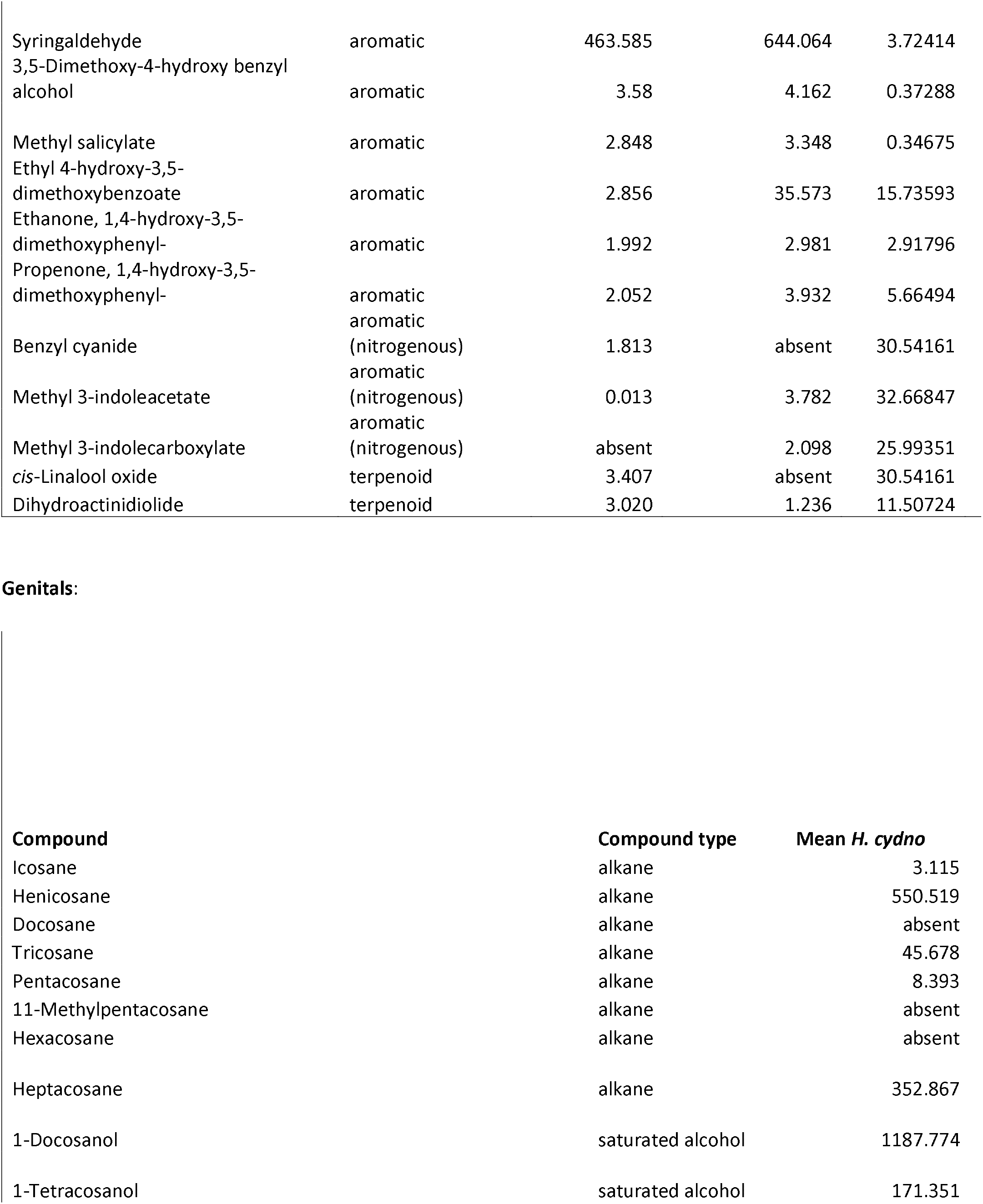
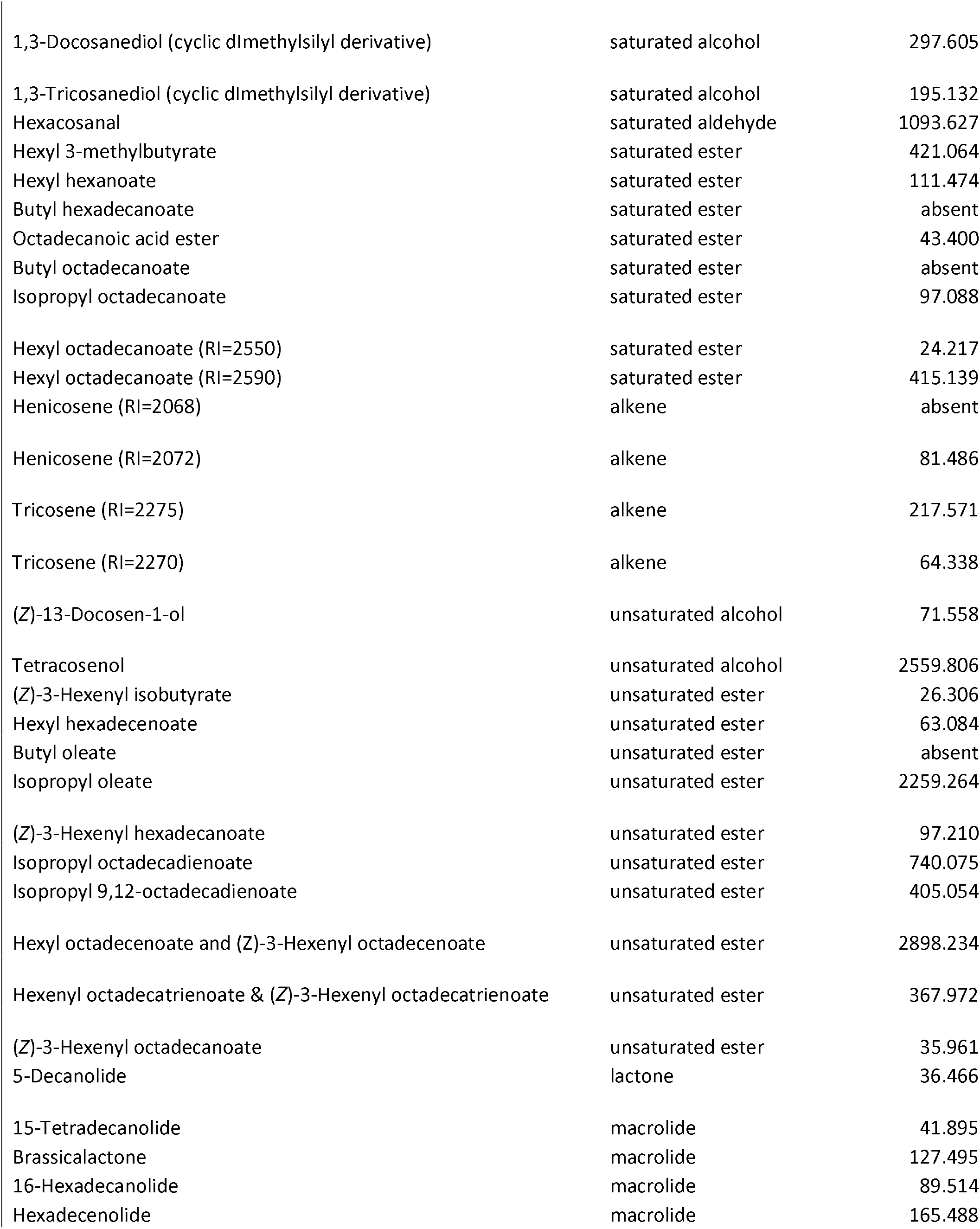
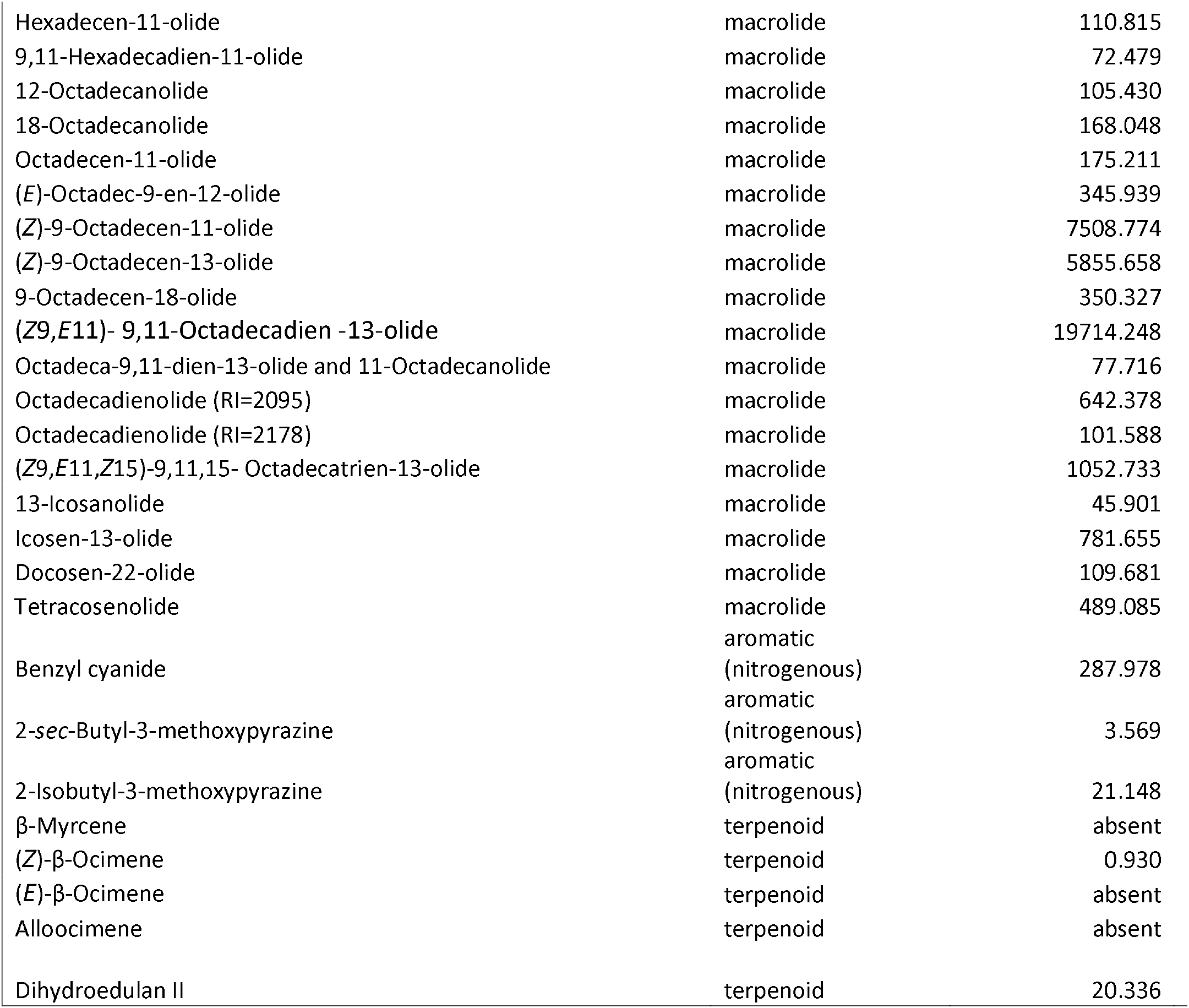
Full list of compounds found in the wings and genitalia of *Heliconius melpomene* and *H. cydno*, details of statistical comparisons of amounts in both species, and cross direction(s) the compound was mapped in, if any. Coefficient of variance in backcrosses is the variance within the backcross direction divided by the mean amount found within that backcross direction, to control for variance differences due to compound abundance differences.

Most compounds mapped did not produce significant QTL at either the p = 0.05 or the Bonferroni-adjusted cutoff levels (the latter taking into account the number of compounds mapped) (Figures S1 and S2). A total of 21 genital compounds and 14 wing compounds produced significant QTL, including three peaks already published in (Byers et al., 2020, Darragh et al., bioRxiv) for octadecanal, 1-octadecanol, and (*E*)-β-ocimene. Of these QTL, nine genital and ten wing compounds were significant at the more conservative Bonferroni threshold (Tables 2 and 3; Table 4). The higher drop-out rate for genital compounds after Bonferroni correction is likely the result of the more stringent correction due to a higher number of compounds corrected for. We also found a single significant QTL on chromosome 18 for relative hindwing androconial area in backcrosses to *H. cydno* (Figure S3). The percentage of parental difference explained by the peak markers at these QTL was highly variable, with values ranging from 0.6-130% for wing compounds and 1.9-170% for genital compounds, but most compounds fell within a range of approximately 20-50% of parental difference explained by the locus. Surprisingly, there was no obvious association between the amount of variance present in the backcrosses and the presence or absence of a QTL for each compound, as seen from the coefficient of variance values in Table 1, which did not differ between compounds that did and did not have a QTL associated (*t* = 0.342, df = 78, p = 0.73).

**Table 2:** Details of compounds with Bonferroni-significant QTL in *Heliconius* wing androconia. a: from (Byers et al., 2020). c: in backcross to *H. cydno*. m: in backcross to *H. melpomene*.

| Compound | Type | Mean (*H. melpomene*, ng)<sup>a</sup> | Mean (*H. cydno*, ng)<sup>a</sup> | Bonferroni-significant QTL on chromosome | QTL position on chromosome (bp) | Percent parental difference explained by genotype at QTL position |
| --- | --- | --- | --- | --- | --- | --- |
| (Z)-9-Octadecenal | Unsaturated aldehyde | 10.94 | Absent | 1<sup>m</sup> | 9955174<sup>m</sup> | 33.544%<sup>m</sup> |
| Benzyl cyanide | Aromatic (nitrogenous) | Absent | 1.813 | 17<sup>m</sup> | 5596551<sup>m</sup> | 67.126%<sup>m</sup> |
| Octadecanal<sup>a</sup> | Saturated aldehyde | 783.4 | 0.569 | 20<sup>c</sup>; 20<sup>m</sup> | 10947362<sup>c</sup>; 11725848<sup>m</sup> | 0.602%<sup>c</sup>; 41.411%<sup>m</sup> |
| Henicosane | Alkane | 12.45 | 1152 | 20<sup>m</sup> | 12126129<sup>m</sup> | 11.291%<sup>m</sup> |
| Methyloctadecanol | Saturated alcohol | 12.23 | Absent | 20<sup>m</sup> | 12374765<sup>m</sup> | 43.435%<sup>m</sup> |
| 1-Octadecanol<sup>a</sup> | Saturated alcohol | 345.6 | Absent | 20<sup>m</sup> | 12652775<sup>m</sup> | 27.709%<sup>m</sup> |
| (Z)-16-Methyl-9-octadecenol | Unsaturated alcohol | 7.501 | Absent | 20<sup>m</sup> | 12652775<sup>m</sup> | 38.888%<sup>m</sup> |
| (Z)-13-Docosenal | Unsaturated aldehyde | 62.56 | 0.276 | 20<sup>c</sup> | 12917911<sup>c</sup> | 28.582%<sup>c</sup> |
| (Z)-11-Icosenal | Unsaturated aldehyde | 150.7 | 3.290 | 20<sup>c</sup> | 13267611<sup>c</sup> | 22.717%<sup>c</sup> |
| Methyl 3-indoleacetate | Aromatic (nitrogenous) | 3.782 | 0.013 | Z<sup>c</sup> | 8478732<sup>c</sup> | 59.220%<sup>c</sup> |

**Table 3:** Details of compounds with Bonferroni-significant QTL in *Heliconius* genitals. a: from (Darragh et al., 2020). c: in backcross to *H. cydno*. m: in backcross to *H. melpomene*.

| Compound | Type | Mean (*H. melpomene*, ng)<sup>a</sup> | Mean (*H. cydno*, ng) | Bonferroni-significant QTL on chromosome | QTL position on chromosome (bp) | Percent parental difference explained by genotype at QTL position |
| --- | --- | --- | --- | --- | --- | --- |
| Hexyl 3-methylbutyrate | Saturated ester | Absent | 421.1 | 3<sup>m</sup> | 234227<sup>m</sup> | 5.863%<sup>m</sup> |
| 2-sec-Butyl-3-methoxypyrazine | Aromatic | 32.66 | 3.569 | 5<sup>c</sup> | 4427722<sup>c</sup> | 38.866%<sup>c</sup> |
|  | (nitrogenous) |  |  |  |  |  |
| 12-Octadecanolide | Macrolide | Absent | 105.4 | 8<sup>c</sup> | 4512378<sup>c</sup> | 31.434%<sup>c</sup> |
| (Z)-9-Octadecen-11-olide | Macrolide | Absent | 7509 | 8<sup>m</sup> | 5341821<sup>m</sup> | 12.192%<sup>m</sup> |
| (Z9,E11)- 9,11-Octadecadien -13-olide | Macrolide | Absent | 19714 | 8<sup>m</sup> | 5341821<sup>m</sup> | 2.826%<sup>m</sup> |
| Hexadecenolide | Macrolide | 6.866 | 165.5 | 8<sup>m</sup> | 5341821<sup>m</sup> | 170.403%<sup>m</sup> |
| Hexyl octadecanoate | Saturated ester | 121.8 | 24.21 | 8<sup>m</sup> | 5987924<sup>m</sup> | 46.335%<sup>m</sup> |
| (Z)-9-Octadecen-13-olide | Macrolide | 7.282 | 5856 | 8<sup>m</sup> | 6072562<sup>m</sup> | 15.975%<sup>m</sup> |
| Isopropyl oleate | Unsaturated ester | 50.22 | 2259 | 14<sup>m</sup> | 7370836<sup>m</sup> | 16.385%<sup>m</sup> |

**Table 4:**
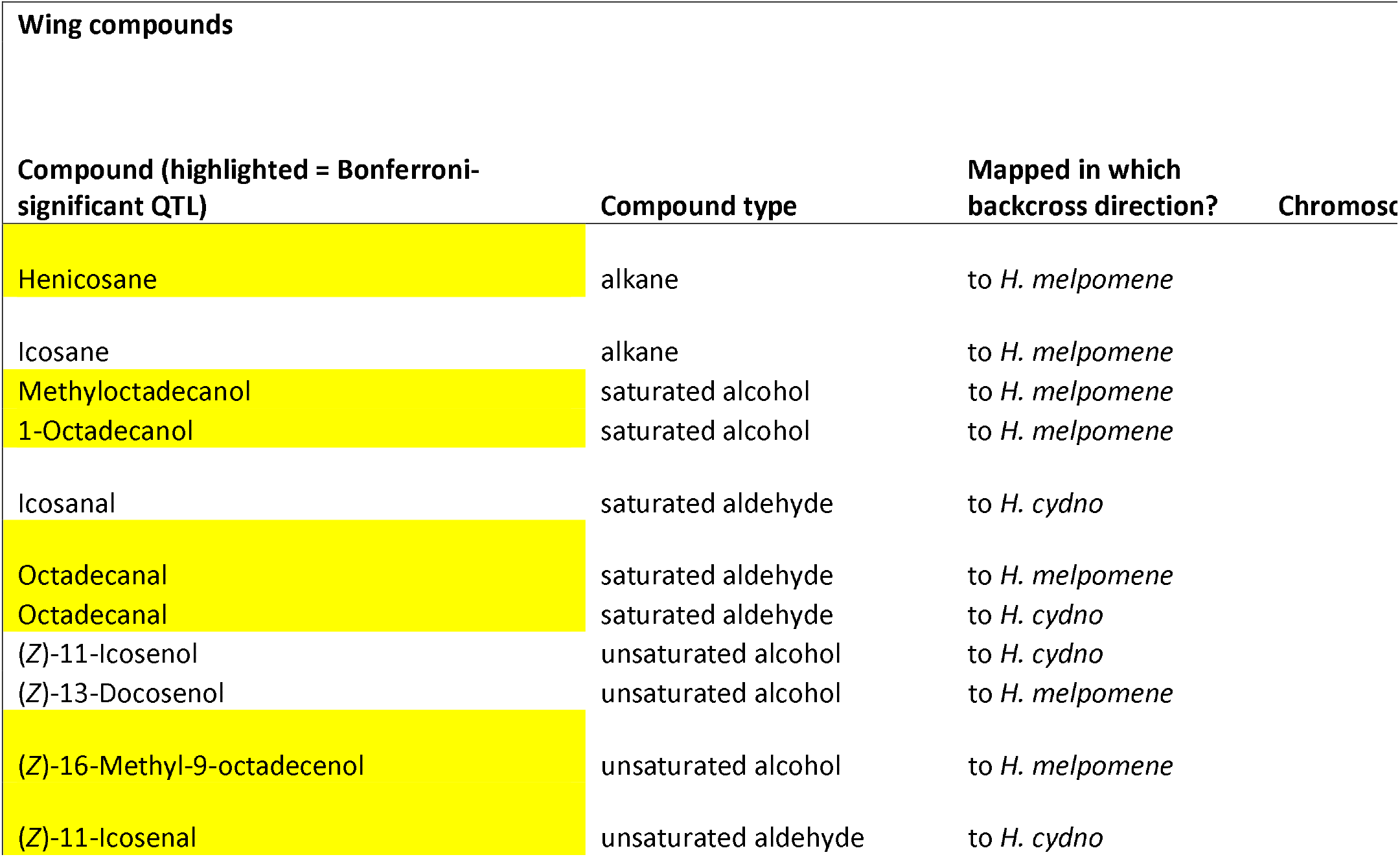
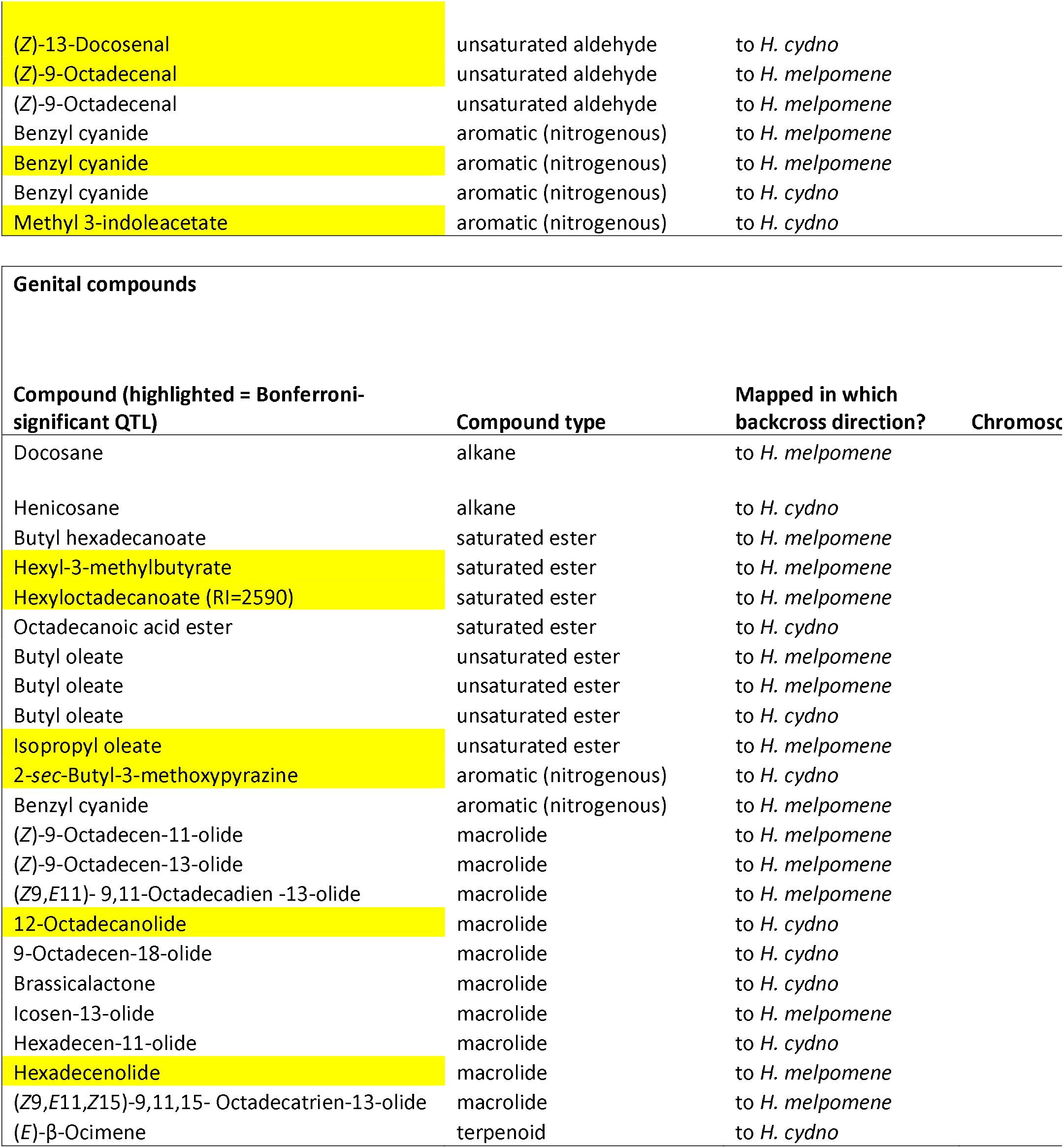
Details of quantitative trait loci passing a significance threshold of either p = 0.05 or a Bonferroni-corrected threshold.

| Wing compounds |  |  |  |
| --- | --- | --- | --- |
| Compound (highlighted = Bonferroni-significant QTL) | Compound type | Mapped in which backcross direction? | Chromosc |
| Henicosane | alkane | to *H. melpomene* |  |
| Icosane | alkane | to *H. melpomene* |  |
| Methyloctadecanol | saturated alcohol | to *H. melpomene* |  |
| 1-Octadecanol | saturated alcohol | to *H. melpomene* |  |
| Icosanal | saturated aldehyde | to *H. cydno* |  |
| Octadecanal | saturated aldehyde | to *H. melpomene* |  |
| Octadecanal | saturated aldehyde | to *H. cydno* |  |
| (Z)-11-Icosenol | unsaturated alcohol | to *H. cydno* |  |
| (Z)-13-Docosenol | unsaturated alcohol | to *H. melpomene* |  |
| (Z)-16-Methyl-9-octadecenol | unsaturated alcohol | to *H. melpomene* |  |
| (Z)-11-Icosenal | unsaturated aldehyde | to *H. cydno* |  |

|  |  |  |
| --- | --- | --- |
| (Z)-13-Docosenal | unsaturated aldehyde | to *H. cydno* |
| (Z)-9-Octadecenal | unsaturated aldehyde | to *H. melpomene* |
| (Z)-9-Octadecenal | unsaturated aldehyde | to *H. melpomene* |
| Benzyl cyanide | aromatic (nitrogenous) | to *H. melpomene* |
| Benzyl cyanide | aromatic (nitrogenous) | to *H. melpomene* |
| Benzyl cyanide | aromatic (nitrogenous) | to *H. cydno* |
| Methyl 3-indoleacetate | aromatic (nitrogenous) | to *H. cydno* |

| Genital compounds |  |  |  |
| --- | --- | --- | --- |
| Compound (highlighted = Bonferroni-significant QTL) | Compound type | Mapped in which backcross direction? | Chromosc |
| Docosane | alkane | to *H. melpomene* |  |
| Henicosane | alkane | to *H. cydno* |  |
| Butyl hexadecanoate | saturated ester | to *H. melpomene* |  |
| Hexyl-3-methylbutyrate | saturated ester | to *H. melpomene* |  |
| Hexyloctadecanoate (RI=2590) | saturated ester | to *H. melpomene* |  |
| Octadecanoic acid ester | saturated ester | to *H. cydno* |  |
| Butyl oleate | unsaturated ester | to *H. melpomene* |  |
| Butyl oleate | unsaturated ester | to *H. melpomene* |  |
| Butyl oleate | unsaturated ester | to *H. cydno* |  |
| Isopropyl oleate | unsaturated ester | to *H. melpomene* |  |
| 2-sec-Butyl-3-methoxypyrazine | aromatic (nitrogenous) | to *H. cydno* |  |
| Benzyl cyanide | aromatic (nitrogenous) | to *H. melpomene* |  |
| (Z)-9-Octadecen-11-olide | macrolide | to *H. melpomene* |  |
| (Z)-9-Octadecen-13-olide | macrolide | to *H. melpomene* |  |
| (Z9,E11)- 9,11-Octadecadien -13-olide | macrolide | to *H. melpomene* |  |
| 12-Octadecanolide | macrolide | to *H. cydno* |  |
| 9-Octadecen-18-olide | macrolide | to *H. cydno* |  |
| Brassicalactone | macrolide | to *H. cydno* |  |
| Icosen-13-olide | macrolide | to *H. melpomene* |  |
| Hexadecen-11-olide | macrolide | to *H. cydno* |  |
| Hexadecenolide | macrolide | to *H. melpomene* |  |
| (Z9,E11,Z15)-9,11,15- Octadecatrien-13-olide | macrolide | to *H. melpomene* |  |
| (E)-β-Ocimene | terpenoid | to *H. cydno* |  |

Compounds that showed significant QTL tended to be of specific chemical classes. For the wings, of the 14 wing compounds with significance at p = 0.05, 12 were fatty acid-derived compounds (FADs), including alkanes as well as oxygenated alkanes and alkenes (aldehydes and alcohols). The remaining two significant wing compounds were both nitrogenous aromatics. For the genitals, 10 compounds producing significant QTL were macrolides and eight were FADs (two alkanes, four saturated esters, and two unsaturated esters). The remaining three compounds were again nitrogenous aromatics (two compounds) and one terpene. This is in general agreement with the distribution of compound types in both body regions. The mapped wing compounds comprise 18 fatty acid-derived compounds, three nitrogenous aromatics, two non-nitrogenous aromatics, and two terpenoids, so the finding of significant QTL mostly for fatty acid-derived compounds is not surprising. For the genitals, the mapped compounds consist of 22 fatty acid-derived compounds, 21 macrolides, four terpenoids, two nitrogenous aromatics, and one lactone.

Significant quantitative trait loci showed some same-chromosome clustering on specific chromosomes (Figure S4). Notably, these clusters did not overlap with chromosomes harboring known QTL for color pattern, mate choice, or other traits, which generally tend not to be clustered in the genome, apart from a close association between mate choice and a wing patterning gene (Figure 3). When either all QTL significant at p < 0.05 or Bonferroni-corrected significant QTL were combined with known QTL, clustering at the chromosomal level was statistically significant (χ^2^ = 64.036 for Bonferroni-significant QTL, χ^2^ =78.524 for all significant QTL, p < 0.001 for both) for chromosomes 8 and 20. In particular, all but one QTL for FAD compounds in the wings mapped to chromosome 20 (the exception mapped to chromosome 1 as well as having a peak on chromosome 20). The other significant wing compounds (all nitrogenous aromatics) mapped to chromosomes 10, 17, 19, and the Z chromosome (chromosome 21). QTL on chromosome 20 were broadly overlapping, in part due to our relatively small mapping populations producing broad Bayesian confidence intervals for QTL location (Figure 4). In addition, the peaks themselves were strongly overlapping on the latter half of chromosome 20. This chromosomal-level clustering was statistically significant (χ^2^ = 178.39 for all significant QTL, χ^2^ = 113.02 for Bonferroni-corrected significant QTL, p < 0.001 in both cases) and residual analysis identified chromosome 20 as the sole outlier. When broken down by chemical class, only FAD compounds clustered significantly at the chromosomal level, again on chromosome 20 (χ^2^ = 232.49 for all significant QTL, p < 0.001), and this was also significant when only Bonferroni-corrected significant QTL were tested (χ^2^ = 137.55, p < 0.001).

**Figure 3:**
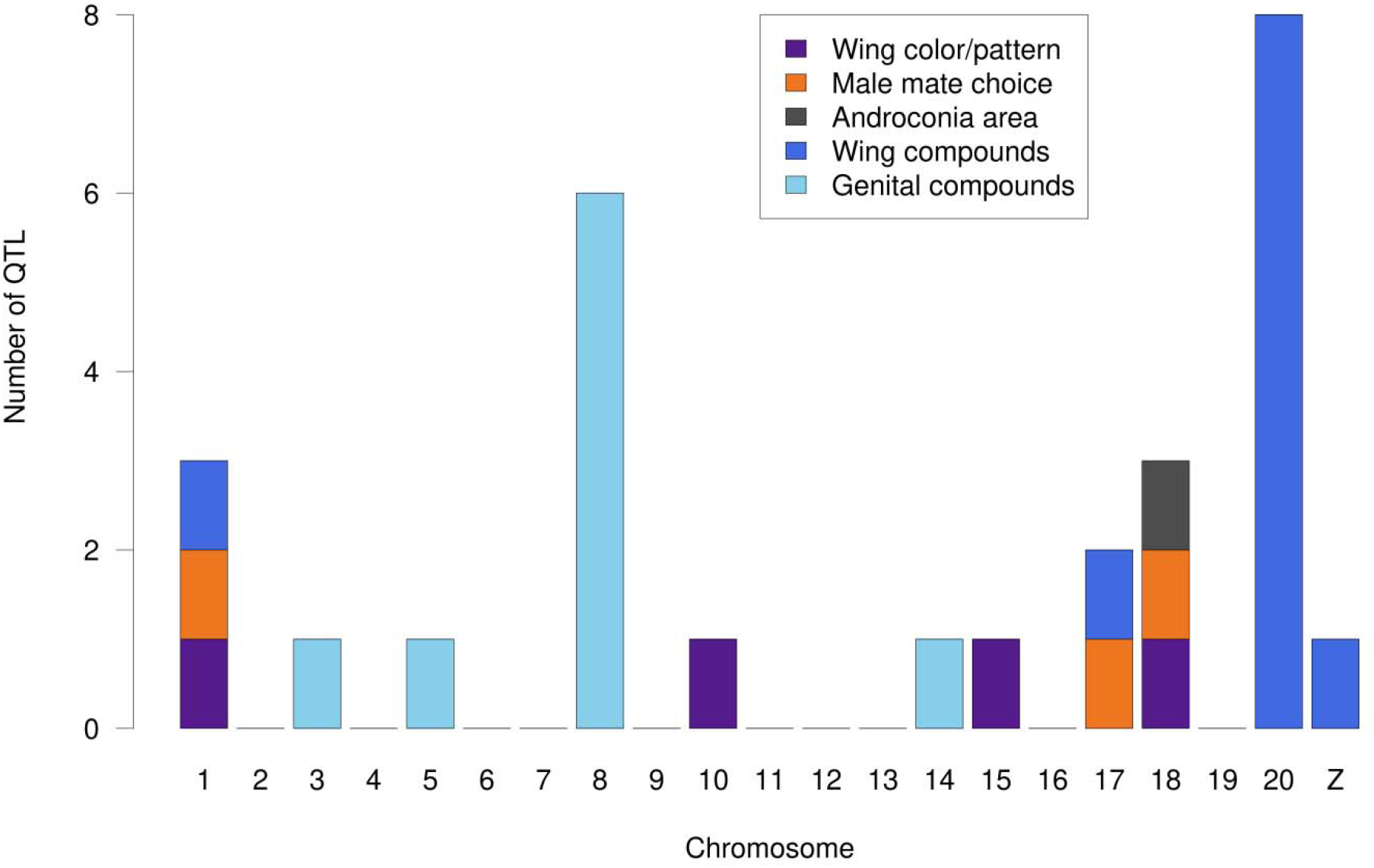
QTL for various traits across the Heliconius melpomene and *H. cydno* genomes. Wing color/pattern loci represent the four major color loci (Jiggins, 2017); male mate choice loci are from (Merrill et al., 2019); (relative) androconia area and Bonferroni-significant wing and genital compound loci from this study.

**Figure 4:**
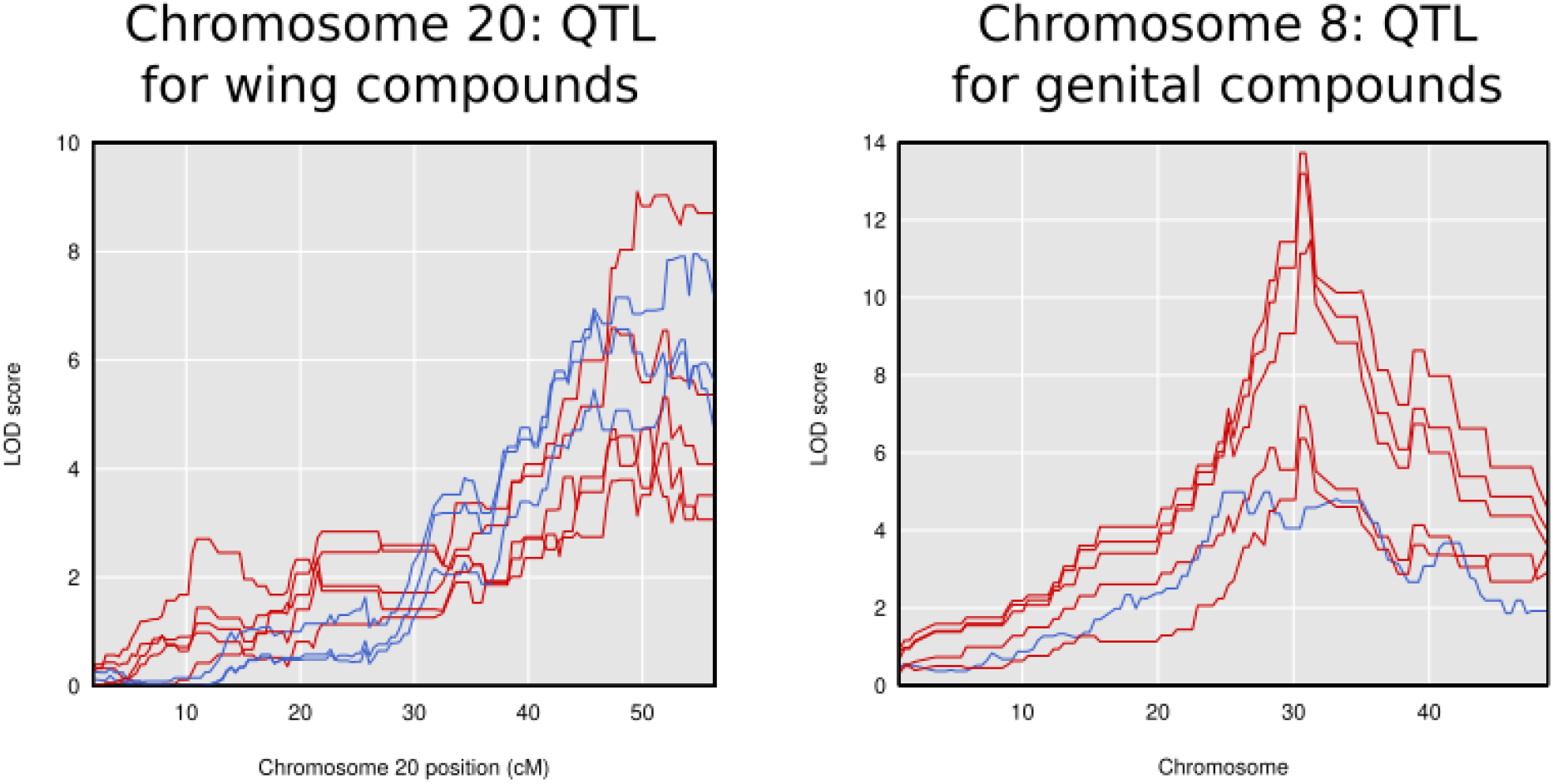
Chromosome-specific QTL plots for compounds with significant QTL (after Bonferroni correction) on those chromosomes, with the entire chromosome length shown. Red: compound mapped in backcrosses to *H. melpomene*; blue: compound mapped in backcrosses to *H. cydno*.

Similar compound class-specific clustering at the chromosome level was seen to a lesser extent for the genitals, where six of the ten macrolides had QTL on chromosome 8, with the others more broadly dispersed on chromosomes 2, 7, 13, 14, and 20. The fatty acid-derived compounds (mostly esters) mapped more broadly across the genome than in the wings, with QTL on chromosomes 1, 2, 3, 6, 8, 9, 14, and 20. Chromosome-level clustering was again statistically significant (χ^2^ = 46.037 for all significant QTL, χ^2^ = 101.83 for Bonferroni-corrected significant QTL, p = 0.0017 and p < 0.001 respectively) and residual analysis identified chromosomes 8 and 14 as the outliers when all QTL were considered, and chromosome 8 when only Bonferroni-corrected QTL were included. The clustering on chromosome 8 is solely due to the presence of macrolide QTL on this chromosome, while chromosome 14 contains QTL for both a macrolide and three FAD compounds. When broken down by chemical class, only macrolide compounds clustered significantly at the chromosomal level in the genitals, again on chromosome 8 (χ^2^ = 58.079 for all significant QTL, p < 0.001), and this was also significant when only Bonferroni-corrected significant QTL were tested (χ^2^ = 110.88, p < 0.001). Again, the QTL peaks themselves were strongly overlapping in the middle of chromosome 8 (Figure 4) rather than being dispersed across the chromosome. Two compounds had significant QTL in both wings and genitals: henicosane and benzyl cyanide. Both data sets showed QTL on the same chromosomes (20 and 17 respectively, with additional QTL for benzyl cyanide in the wings on other chromosomes). The henicosane QTL were within 1.5cM of one another, while the benzyl cyanide QTL on chromosome 17 were 2.3cM apart.

The strong clustering of QTL for wing fatty acid-derived compounds on chromosome 20 suggested a common gene or set of linked genes responsible for fatty acid-derived compound metabolism or its regulation. We searched the region between 43.11 - 56.37 cM (9,398,348 - 14,585,564 basepairs) for potential candidate genes that could be responsible for this clustering. This region spanned the confidence intervals for all Bonferroni-significant QTL apart from 1-octadecanol and contained 363 genes. Annotated proteins from this region (from Lepbase, Challis et al., 2016) were searched using BLASTp against the NCBI non-redundant (nr) database, revealing a total of 20 genes potentially involved in the biosynthesis of the wing compounds within this confidence interval on chromosome 20. These genes include two acetyl-CoA carboxylases (involved in primary metabolism), fifteen fatty acyl-CoA reductases (FARs), and two alcohol dehydrogenases (the latter two families involved in both primary and secondary metabolism), none of which have been functionally characterized. Comparing our candidates to those identified in (Liénard et al., 2014), none of our FARs fall within the pheromone gland FAR (pgFAR) clade, whose members fall on chromosome 19. Inspecting the genome for additional secondary metabolism fatty acid-derived metabolism genes, we identified clusters of FARs on chromosomes 19 and 20 (Figure 5), with a more even distribution of alcohol dehydrogenases and desaturases across the genome. Notably, we did not identify any QTL on chromosome 19 for fatty acid-derived compound production in the wings.

**Figure 5:**
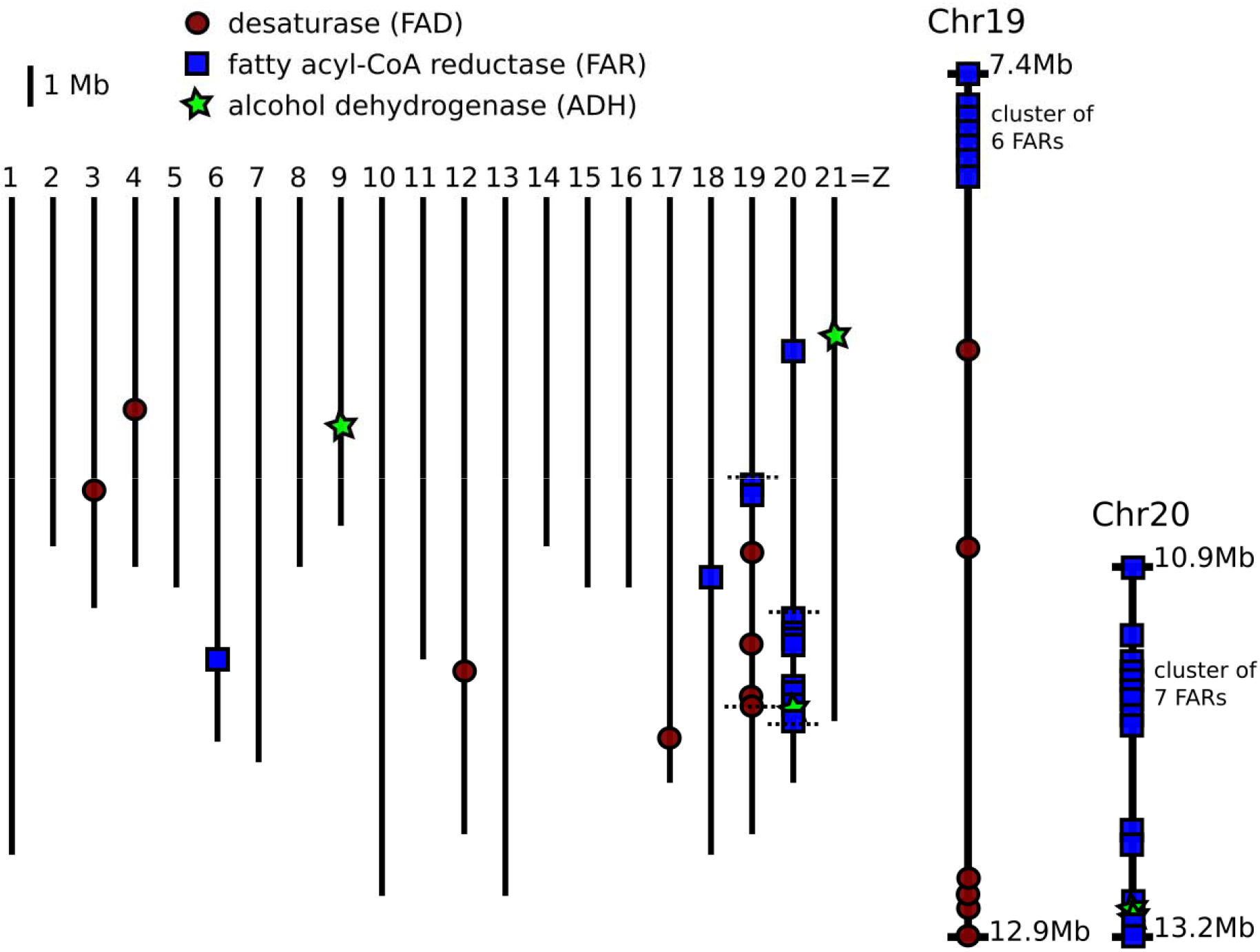
Location of candidate genes for fatty acid-derived compound biosynthesis across the genome of *Heliconius melpomene*. Horizontal dotted lines on chromosomes 19 and 20 indicate the endpoints of zoomed in regions.

## Discussion

The genetic basis for traits contributing to reproductive isolation remains poorly studied. This is especially the case for differences in chemical signaling between closely related non-model species, in particular for species with complex chemical signatures where multiple components may play a role in mate choice or species recognition. Here, we mapped a total of 40 QTL controlling the segregation of wing and genital pheromone bouquet compounds in crosses between *Heliconius melpomene* and its close relative *H. cydno*. There was significant evidence for non-random clustering of QTL, most notably for wing compounds on chromosome 20, which showed broadly overlapping QTL with closely spaced peaks. There could be a number of reasons for this clustering of loci. First, theory predicts that mutations controlling species differences may accumulate, and persist, in regions already resistant to gene flow, such that recombination is less likely to disrupt species differences (Smadja & Butlin, 2011). Second, functionally similar genes are likely to occur in close linkage due to *e.g.* tandem gene duplication, such that chemically similar compounds could be regulated by linked genes due to functional similarities between their synthesis pathways (Osbourn, 2010). Finally, related compounds could differ between species as a pleiotropic effect of the same mutation if they are downstream products of the same biochemical pathway or in the case of a multifunctional enzyme. The end result of all of these processes is linkage that can facilitate speciation by impeding recombination that will otherwise disrupt the genetic associations between traits that characterize emerging species.

In the case of the compounds described here, it is likely that shared biosynthetic origins play an important role in their chromosomal distributions. The majority of the wing compounds found in either species are fatty acid-derived compounds (FADs; 19 of 31), and the majority of these FADs (12 of 19) were associated with a significant QTL on chromosome 20. It is interesting that there was clustering of one class of biosynthetic enzyme in this pathway, the fatty acyl-CoA reductases (FARs), on both chromosomes 19 and 20, but no QTL on chromosome 19 for any wing or genital FAD compound, suggesting the FARs on chromosome 19 are not involved in putative pheromone compound production. Similarly, QTL for genital compounds exhibited slight clustering of loci underlying macrolide compound production on chromosome 8, but otherwise were more evenly spread across the genome, likely due to their greater biosynthetic diversity. The QTL for shared macrolide compounds may also be due to shared biosynthetic origins of these compounds, but the biosynthesis of macrolides is less well understood and consequently we were unable to identify candidate genes on this chromosome. We also found a QTL on chromosome 3 for hexyl 3-methylbutyrate, a known bioactive component of the genital pheromone of *H. cydno* (Estrada, 2009) whose genetic basis was previously unknown. Of note, the QTL for relative androconial area roughly peaks on the gene *optix* (chromosome 18), a locus controlling red color patterns in *Heliconius* (Reed et al., 2011, Van Belleghem et al., 2017). This locus is also known to control male-specific scales in the basal *heliconiine Dryas iuila* and has an expression domain in the hindwing androconia region (Martin et al., 2014). This overlap raises the possibility for coupled control of color pattern and androconial pheromone production differences between *Heliconius* species.

Overall, we found no evidence for linkage of any of the chemical QTL clusters with other loci or QTL that have been shown to play a difference in the strong assortative mating observed between *H. melpomene* and *H. cydno*. The major wing patterning genes are located on chromosomes 1 (*aristalless*), 10 (*WntA*), 15 (*Cortex*), and 18 (*optix*), only chromosome 1 of which harbors a chemical QTL, but the peak does not include the *aristalless* gene. In addition, major QTL for male mate choice map to chromosome 1, 17, and 18 (Merrill et al., 2019), but, similarly, the chemical QTL we identified on chromosomes 1 and 17 do not overlap with the major QTL identified for male mate choice. We might expect an overlap between chemical QTL and female mate choice QTL (as females likely assess males partially based on their wing androconial chemistry), but the latter has not been mapped in this species pair. Nonetheless, the clustering of wing compounds in particular might contribute to pheromone-based speciation, particularly in the face of gene flow (Via, 2012). If more than one pheromone component that maps to chromosome 20 is involved in reproductive isolation, and if genes responsible for their production are tightly linked (or if pleiotropy occurs), this may also facilitate speciation (Felsenstein, 1981, Smadja & Butlin, 2011, Merrill et al., 2019) and prevent the breakup of potentially adaptive pheromone bouquet mixtures. Evidence suggests that mixture effects are important in pheromone processing (Riffell et al., 2009, Clifford & Riffell, 2013, Lei et al., 2013), and these mixtures might be maintained through tight genetic linkage between biosynthetic loci, facilitated by gene evolutionary events such as tandem duplication. In contrast to the tight clustering seen on chromosome 20 for wing FAD compounds, we saw more limited clustering for the more chemically diverse genital compounds, likely due to their more diverse potential biosynthesis pathways. This might hamper any potential role in reproductive isolation. Given their known function as anti-aphrodisiacs, genital compounds are not so likely to play a major role in isolation between *Heliconius* species (Gilbert, 1976, Schulz et al., 2007, Estrada, 2009).

Some of the compounds found on wings and in genitalia might have a plant origin. Several compounds in the wings and genitals did not differ between parental species, and many of these have been shown to be affected by larval diet in *Heliconius melpomene*. Most of the invariant wing compounds are aromatics, which are expected to be plant derived in *Heliconius* (Darragh et al., 2019). Notably, we reared all mapping butterflies on the same larval and adult diets, thus effectively creating a dietary common garden experiment, and so we do not expect larval or adult diet to affect the result of the QTL mapping. Many male moth pheromone components are derived from host plant volatiles with minimal alterations, potentially due to females’ pre-existing biases for detecting these compounds during the host plant search (Conner & Iyengar, 2016). However, many of our mapped *Heliconius* compounds do appear to have a genetic basis (i.e. a QTL was found), in line with the biosynthesis of FAD and related compounds by female moths. Breaking our compounds down into four classes (FADs, macrolides, aromatics, and terpenoids), we failed to find QTL (at the p = 0.05 level) for 54%, 58%, 43%, and 83% of these classes respectively across both wings and genitals. In particular, terpenoids that differ between species may be the result of differential larval host plant use. It is of course possible that plant derived compounds could show a genetic basis if there were species differences in the genes regulating their uptake, and one example might be icosanal, which has been shown to be influenced by larval host plant (Darragh et al., 2019) but which is also associated with a QTL in this study.

This study provides a first step towards better characterizing the genes causing chemical differences between species. Fatty acid-derived compound biosynthesis candidate genes and QTL for FAD compounds are both clustered on chromosome 20. These FAD compounds are produced from acetyl-CoA followed by rounds of chain elongation, desaturation, fatty acyl-CoA reduction, oxidization, and acetyltransferase activity to produce the final pheromone compounds. The fatty acyl-CoA reduction step, key to pheromone biosynthesis, uses FARs to remove the CoA moiety and return an alcohol. More than half of these FAR candidates are found in tandem duplications, as are the two alcohol dehydrogenase candidates. These duplications could have provided raw material for the origin of new biosynthetic function and thus increased pheromone diversity. FARs also cluster on chromosome 19, and these FARs (unlike those on chromosome 20) fall within the pheromone gland FAR (pgFAR) subfamily (Liénard et al., 2014, Löfstedt et al., 2016). However, these phylogenetic analyses of FARs in *Heliconius, Bombyx, Danaus*, and *Bicyclus* do not generally show lineage-specific clustering of the chromosome 20 FARs; instead, they are dispersed throughout the tree. This suggests an ancient evolutionary origin of the *Heliconius* chromosome 20 FAR cluster, in contrast to *Heliconius*-specific duplications of chromosome 19 FARs in the pgFAR subfamily which are clustered in the tree.

Single FAR and alcohol dehydrogenase genes are missing from *H. cydno* (Byers et al., 2020), suggesting an obvious mechanism for the striking difference in FAD compounds between the two species; these differences may also be due to e.g. differential regulation of FARs or other biosynthetic genes, or mutations within these genes. Alternately, a single FAR gene or small set of genes may be responsible for the biosynthesis of the majority of FAD compounds in *Heliconius*, similar to the situation found with *eloF* in *Drosophila*, where knockdown of the gene produces widely pleiotropic effects on cuticular hydrocarbon profiles (Combs et al., 2018). Similar effects are seen in *Spodoptera litura*, where RNAi of *FAR17* alone alters levels of the four major sex pheromone components (Lin et al., 2017). Another option, though one we consider unlikely, is that this tight cluster of FARs on chromosome 20 serves as a supergene, allowing multiple phenotypic traits (here individual chemical compounds) to co-segregate and maintaining strong integration of the chemical phenotype. Supergenes have been described for other phenotypic morphs in *Heliconius* (Joron et al., 2011), but we are unaware of mechanisms on chromosome 20 that might contribute to supergene formation and maintenance, for example a chromosomal inversion, which would have likely been evident in our linkage map.

Information on potential biosynthetic gene clustering in other Lepidoptera species is largely lacking, as most studies have produced transcriptomes of pheromone glands rather than linkage maps or transcriptomes mapped to genomes. Additionally, as much work in Lepidoptera has focused on single pheromone components and thus single biosynthetic genes (Groot et al., 2016), clustering of QTL and candidate genes is rarely discussed. The lack of annotated genomes in many lepidopteran species also can prevent in-depth assessment of clustering of candidate genes, requiring mapping QTL linkage groups back to *Bombyx* or other well-annotated genomes. However, such clustering has been seen in *Nasonia* wasps, where both QTL for multiple cuticular hydrocarbon components and candidate desaturase genes overlap strongly across the genome (Niehuis et al., 2011, 2013). Clusters of acetyltransferases and desaturases exist in *Heliothis* (inferred from synteny with *Bombyx mori*), but the acetyltransferases do not appear to underlie the QTL for sex pheromone differences between *H. virescens* and *H. subflexa* (Groot et al., 2013, 2014). Quantitative trait loci for multiple compounds were investigated in *Heliothis*, finding multiple chromosomes responsible for the production of the same compound classes (all FADs), with more limited clustering (Sheck et al., 2006). This is similar in pattern to what we found in the genital pheromone components, although the latter cover multiple compound classes.

Understanding the genetic architecture of complex traits can allow us to understand their potential role in speciation. Determining whether the observed clustering QTL on chromosomes 8 and 20 reflect tight linkage of multiple biosynthetic and/or regulatory genes or a single “master” locus responsible for the biosynthesis or regulation of multiple pheromone components will contribute to our understanding of both pheromone biosynthesis and pleiotropy and linkage in general. With genes in hand, CRISPR (a technique that works in *Heliconius*, Livraghi et al., 2018) could be used to alter gene function, allowing confirmation of the role of pheromone components via behavioral assays. As major effect loci (as these QTL appear to be) and tight linkage (as potentially seen here) can both facilitate the speciation process, especially in the face of gene flow (Felsenstein, 1981, Merrill et al., 2010, Smadja & Butlin, 2011, Via, 2012), the QTL identified here could be barrier or speciation genes in *Heliconius*. These specific QTL may be involved in interspecific mate choice and reproductive isolation, and thus the genes underlying them could contribute to the maintenance of species boundaries. We have here identified a total of 40 QTL underlying species differences in 33 potential pheromone components, demonstrated clustering of these QTL for wing and genital compounds on specific chromosomes not linked to known loci for species differences, and identified candidate genes underlying the production of the major chemical class of wing compounds. Together, these findings further our knowledge of chemical ecology, pheromone genetics, and their potential roles in *Heliconius* butterfly mate choice and speciation.

## Acknowledgments

Permits for research and collection of butterfly stocks were provided by the government of Panama. KJRPB, KD, IAW, and CDJ were funded by the European Research Council (FP7-IDEAS-ERC 339873); KD was additionally funded by a Natural Environment Research Council Doctoral Training Partnership (Grant No. NE/L002507/1) and a Smithsonian Tropical Research Institute Short Term Fellowship; PMAR was funded by the Jane and Aatos Erkko Foundation; WOM was funded by the Smithsonian Tropical Research Institute; and SS was funded by the Deutsche Forschungsgemeinschaft (grant DFG Schu984/13-1).

## Appendix: Supplemental figures

**Supplemental Figure 1:**
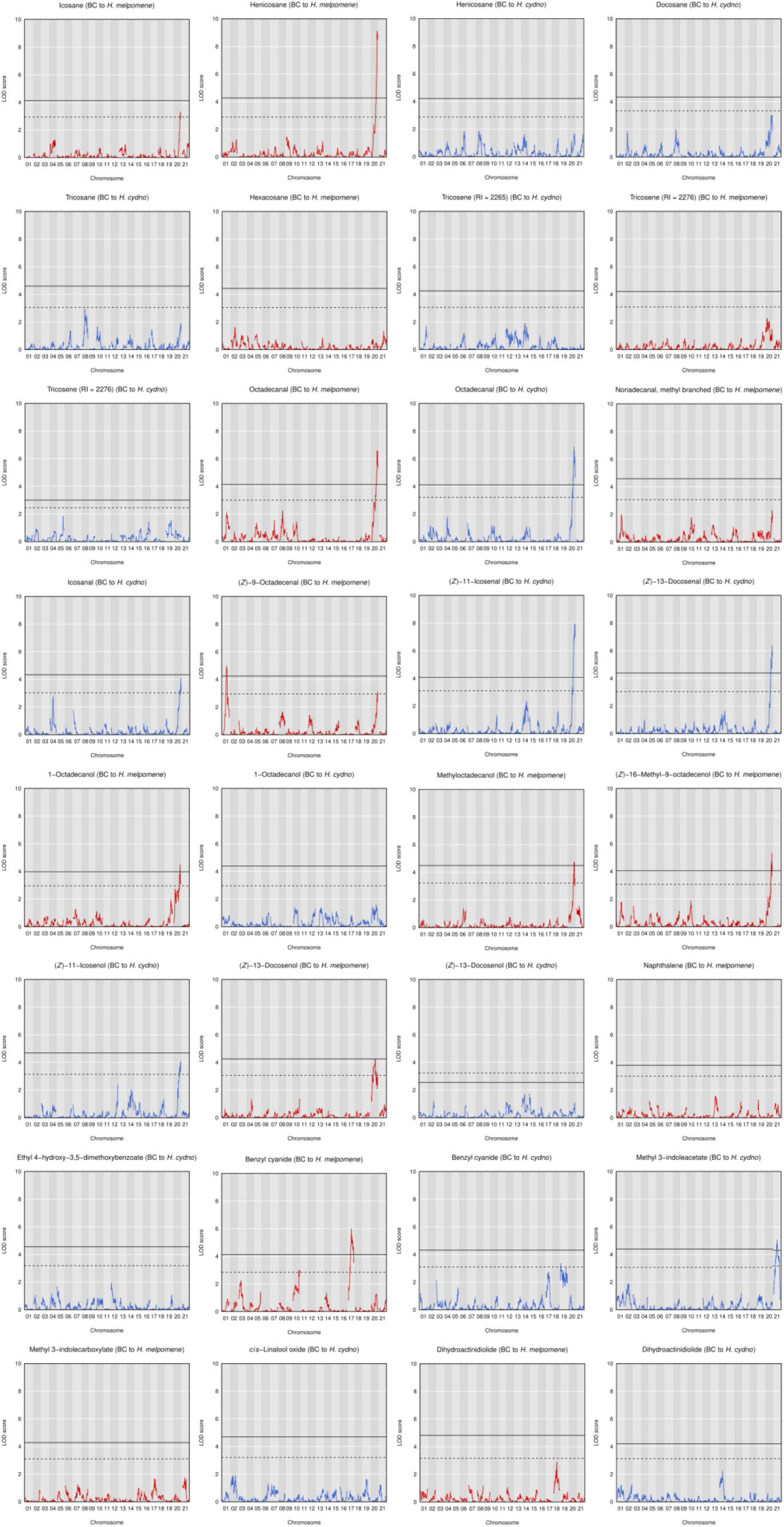
QTL plots for all mapped wing compounds. Dashed horizontal line: p = 0.05 significance threshold. Solid horizontal line: Bonferroni-corrected significance threshold. BC, backcross.

**Supplemental Figure 2 (in two parts):**
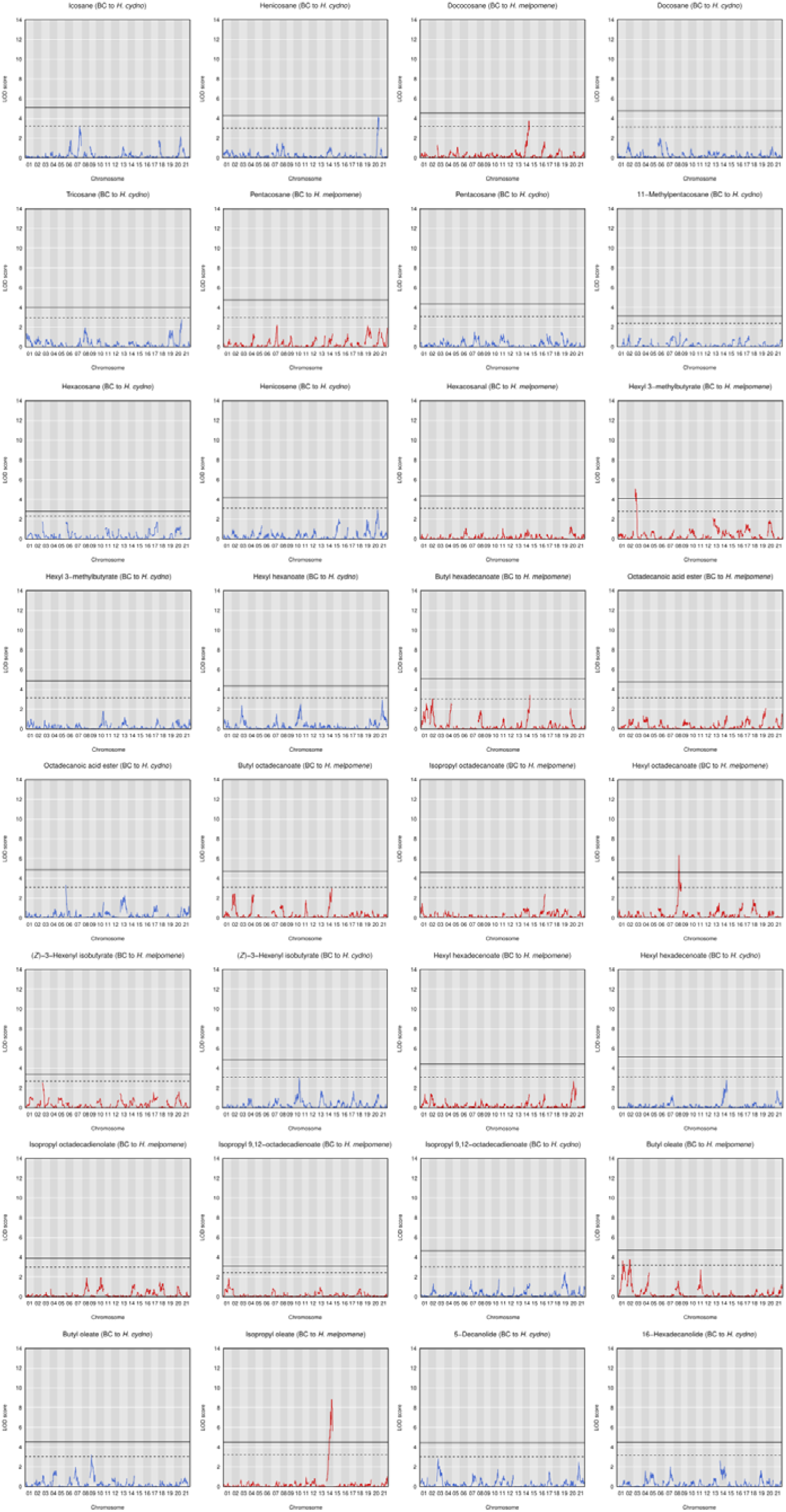
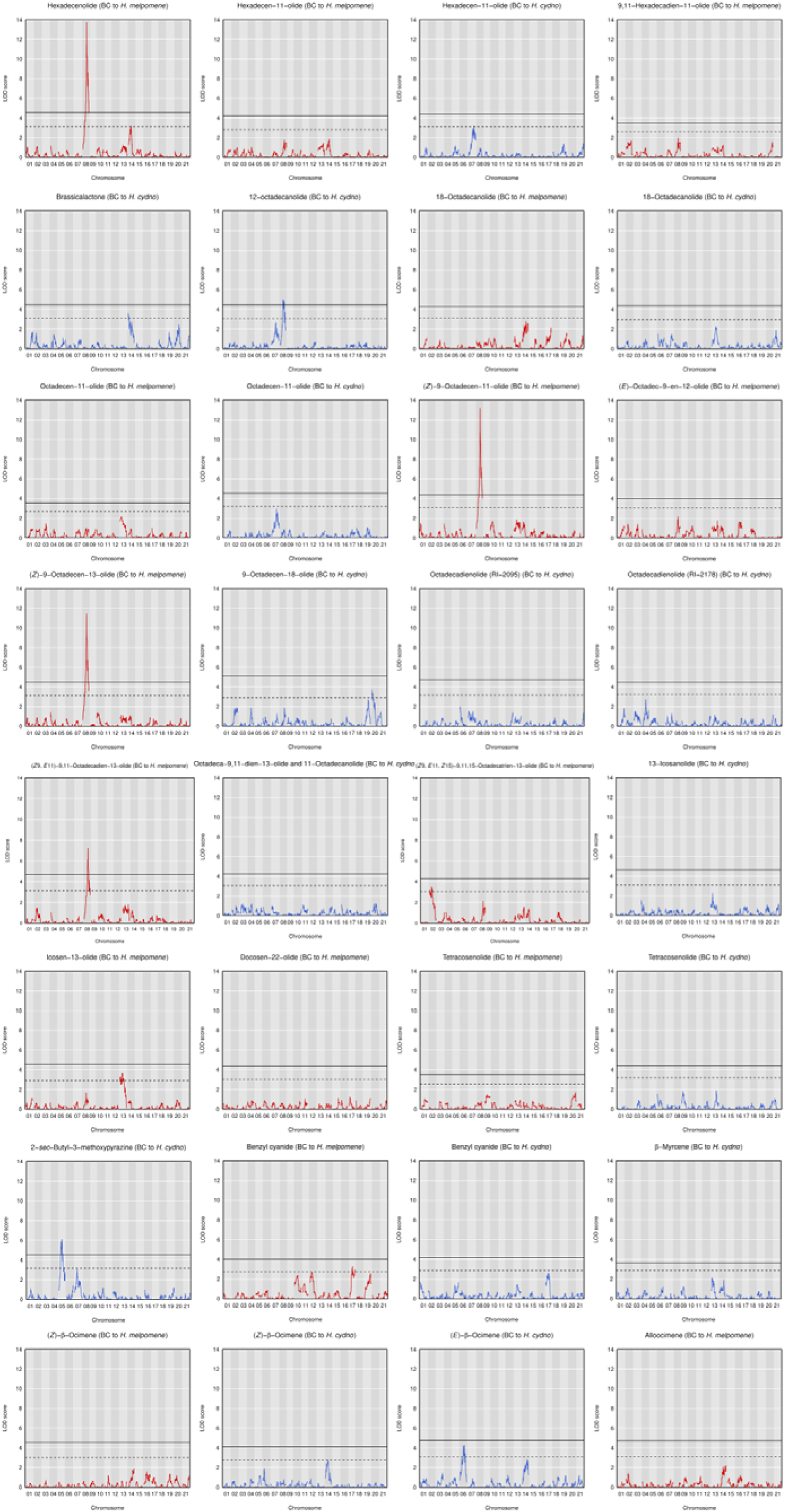
QTL plots for all mapped genital compounds. Dashed horizontal line: p = 0.05 significance threshold. Solid horizontal line: Bonferroni-corrected significance threshold. BC, backcross.

**Supplemental Figure 3:**
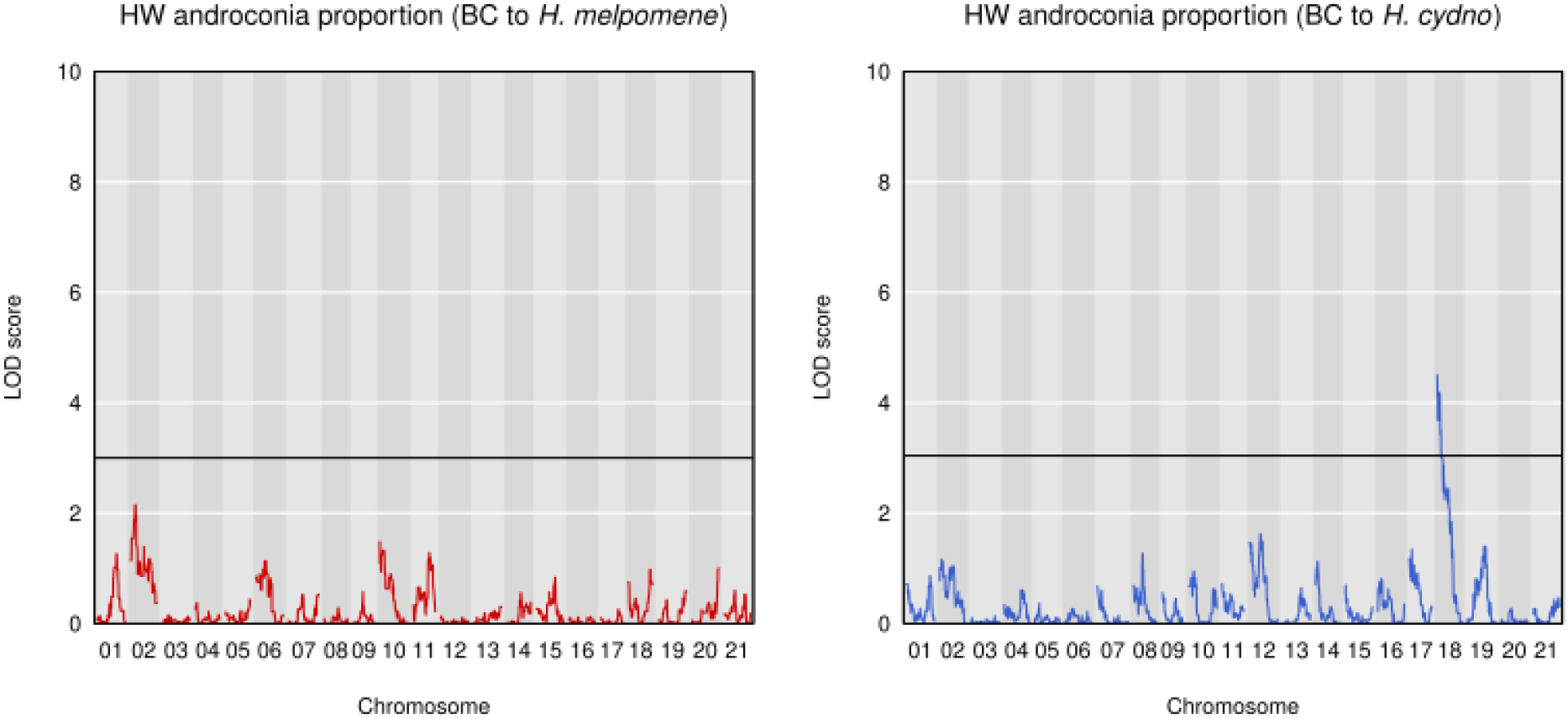
QTL plots for hindwing androconial proportion. Solid horizontal line: p = 0.05 significance threshold. HW, hindwing. BC, backcross.

**Supplemental Figure 4:**
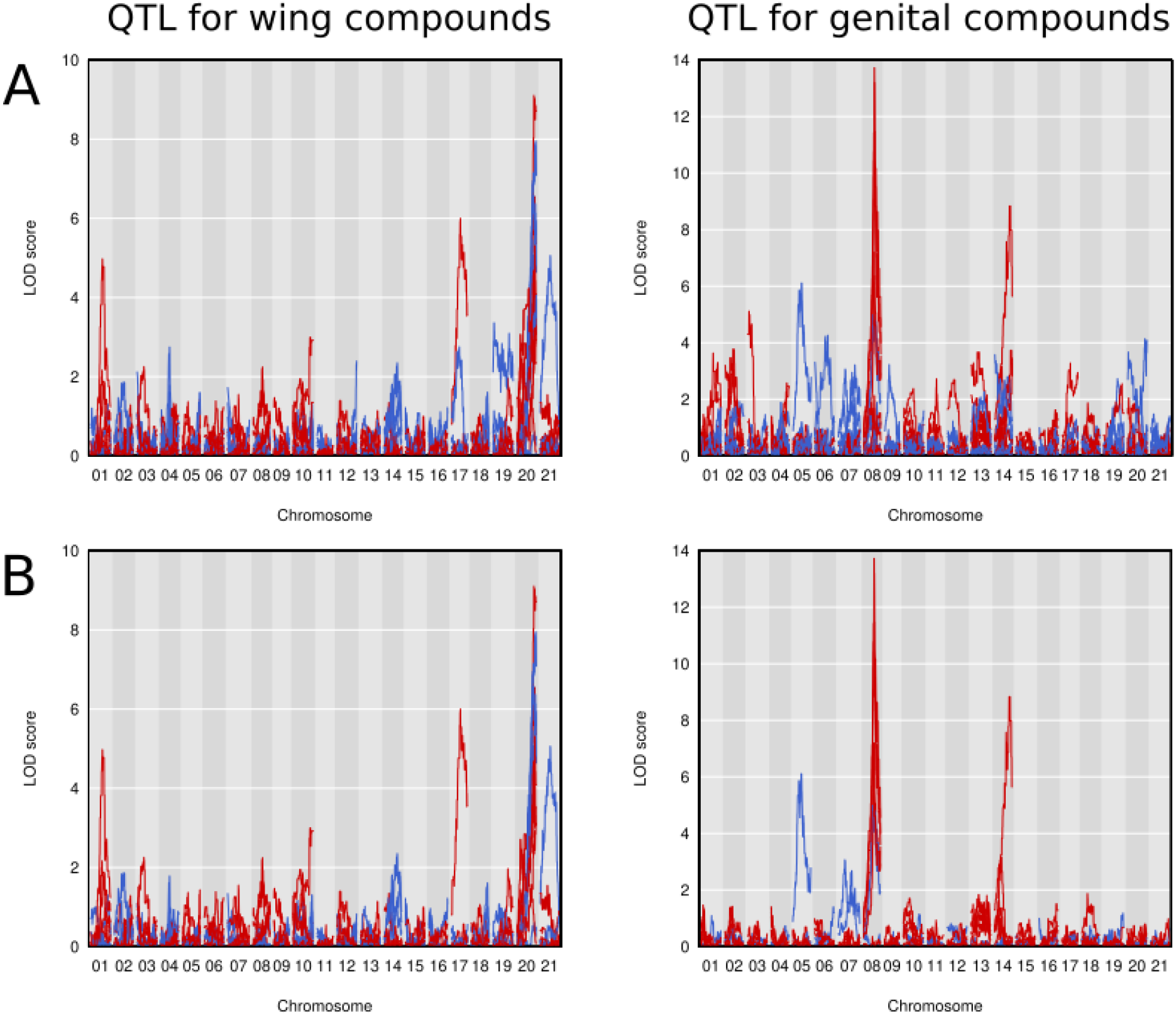
Whole-genome QTL plots for compounds with at least one significant QTL. Red: compound mapped in backcrosses to *H. melpomene*; blue: compound mapped in backcrosses to *H. cydno*. A: Compounds with at least one QTL significant at the p = 0.05 level shown. B: Compounds with at least one QTL significant after Bonferroni correction shown.

